# Inhibitory effects of Akt on neuronal FoxO4 activation after subarachnoid hemorrhage *in vivo* and *in vitro*

**DOI:** 10.1101/2020.01.20.912436

**Authors:** Min Qi, Sheng-qing Gao, Jia-qiang Liu, Yan-ling Han, Bin Yuan, Xiao-ming Zhou, Xi-lin Liu, Meng-liang Zhou, Lan-rong Zheng

## Abstract

Secondary brain injury following subarachnoid hemorrhage (SAH) is the critical contributor to the mortality of SAH patients. The underlying mechanisms are poorly understood. In this study, we utilized a mice model of SAH to investigate whether FoxO4 is related to the brain injury after SAH and identified its upstream regulator Akt. Experimental SAH was induced in adult male mice by prechiasmatic cistern injection. Brain FoxO4 protein levels in cytoplasm and nucleaus were examined in the sham-operated controls, and in mice 1h, 6h, 12h, 24h, 3d, and 5d after SAH induction. The Akt inhibitor LY294002 was administered by intracerebroventricular infusion to determine its effects on FoxO4. Moreover, the expression of FoxO4 was also investigated in neurons incubated with hemoglobin *in vitro*, which was also dertermined after inhibition of Akt. FoxO4 protein expression in the nuclei increased remarkably after SAH. The Akt inhibitor LY294002 induced more FoxO4 nuclear localization after SAH *in vivo* and *in vitro*. Our results suggest the activation of FoxO4 after SAH and which was inhibited by the increased phosphorylated Akt (p-Akt).

## 1. Introduction

Subarachnoid hemorrhage (SAH) is the main type of stroke which account for about 20% in total in developed countries [1]. SAH has attracted much attention in the past decades for the high morbidity and mortality. About 25% of patients die almost in 72 hours after hemorrhage. The primary cause of mortality in SAH patients is early brain injury, that is, the pathological processes in the first 72 hours after initial bleeding. However, few treatments are available for early brain injury because its underlying mechanisms remain unclear till now [2].

Accumulating evidence suggests that apoptosis is involved in early brain injury after SAH. Apoptotic neuronal death could be related to the high morbidity and mortality in SAH patients [3]. The serine-threonine kinase, Akt, was found to play a crucial role in the apoptosis in neurons after SAH [4]. The Akt was phosphorylated after activcation, inhibiting the downstream molecules to make neurons become resistant to apotpotic stimuli. In an animal model of SAH, it was reported that glycogen synthase kinase-3β (GSK3β) is the one regulated by Akt 4. Besides GSK3β, FoxO4 was also found to be induced nuclear exclusion after Akt phosphorylated in cancer and some other diseases [5–7].

FoxO proteins are a family of transcription factors with four members in mammals, namely FoxO1, FoxO3a, FoxO4, and FoxO6. They are originally identified as downstream regulators of the insulin pathway, are known to bind to the promoters of a broad variety of target genes and be involved in diverse cellular and physiological processes including cell proliferation, apoptosis, reactive oxygen species (ROS) response, longevity, cancer and regulation of cell cycle and metabolism [8]. Sereval studies indicated that FoxO4 seems to be more important to the central nervous system(CNS) injuries or diseases [9,10]. Hence, we investigated the expression and activation of FoxO4 and the underlying regulatory role of Akt to activation of FoxO4 after SAH.

## 2. Results

### 2.1. The expressions and locations of FoxO4 and p-Akt in the brain after SAH in vivo

By Western blotting, we analyzed the FoxO4 and p-Akt expressions in the brain. FoxO4 was not changed significantly in the total protein (Fig.1B, F). It was found to be decreased in the nucleus im mediately at 1h after SAH and then increased at 6h, 12h and 24h. It was decreased again at 3d and 5d(Fig.1A, E). In the cytoplasmic protein, the levels of FoxO4 were increased from 6h, peaked at 12h(Fig.1C, G). These findings showed the nuclear translocation of FoxO4 after SAH.

**Figure 1.**
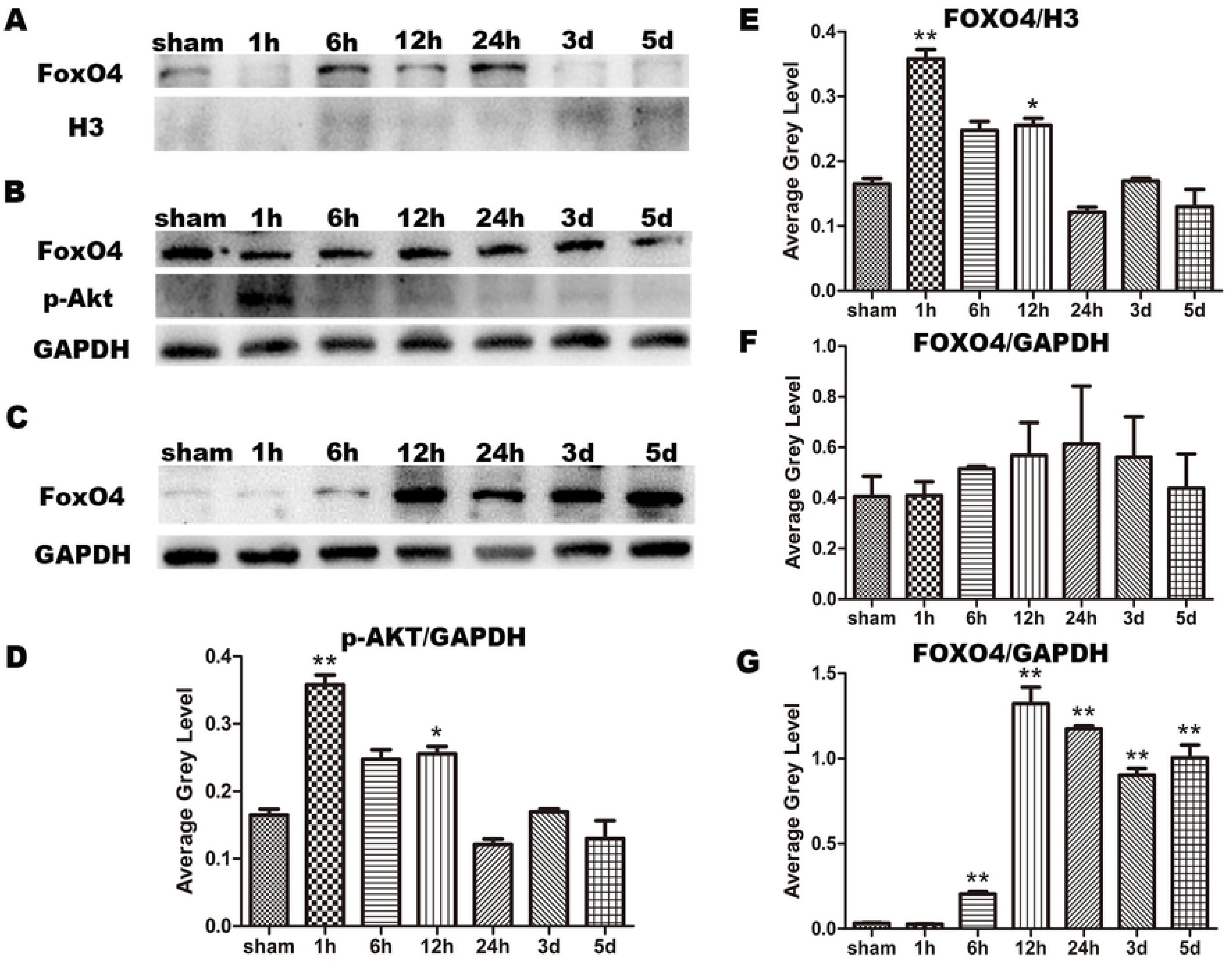
The expression of FoxO4 and p-Akt in the brain. The nucleoprotein content of FoxO4 at different time periods after SAH was detected (A). Western blotting was used to detect the total protein content of FoxO4 and p-Akt at different time periods after SAH (B). In cytoplasmic protein, the content of FoxO4 in different time periods after SAH was detected (C).Statistical analysis of p-Akt changes at different time periods, n = 3, \**P* < 0.05 vs. sham, \*\**P* <0.01 vs. sham(D). Statistical analysis of nucleoprotein content of FoxO4 at different time periods, n = 3, \**P* < 0.05 vs. sham, \*\**P* < 0.01 vs. sham (E).Statistical analysis of the total protein content of FoxO4 changes at different time periods, n = 3, \**P* < 0.05 vs. sham (F).Statistical analysis of the cytoplasmic protein content of FoxO4 changes at different time periods, n = 3, \**P* < 0.05 vs. sham, \*\**P* < 0.01 vs. sham (G).

For p-Akt, it was found to be increased from 1h to 12h after SAH (Fig1 B, D). Because of the early activation of Akt, we treated the animals with Akt inhibitor LY294002 when SAH was induced.At 1h after SAH, we could find the p-Akt positive fluorescence in cytoplasm of neurons(Fig. 2A, B). While stronger fluorescence for FoxO4 was found in the neuronal cytoplasm and nucleus after SAH compared with the sham group (Fig.2C, D).

**Figure 2.**
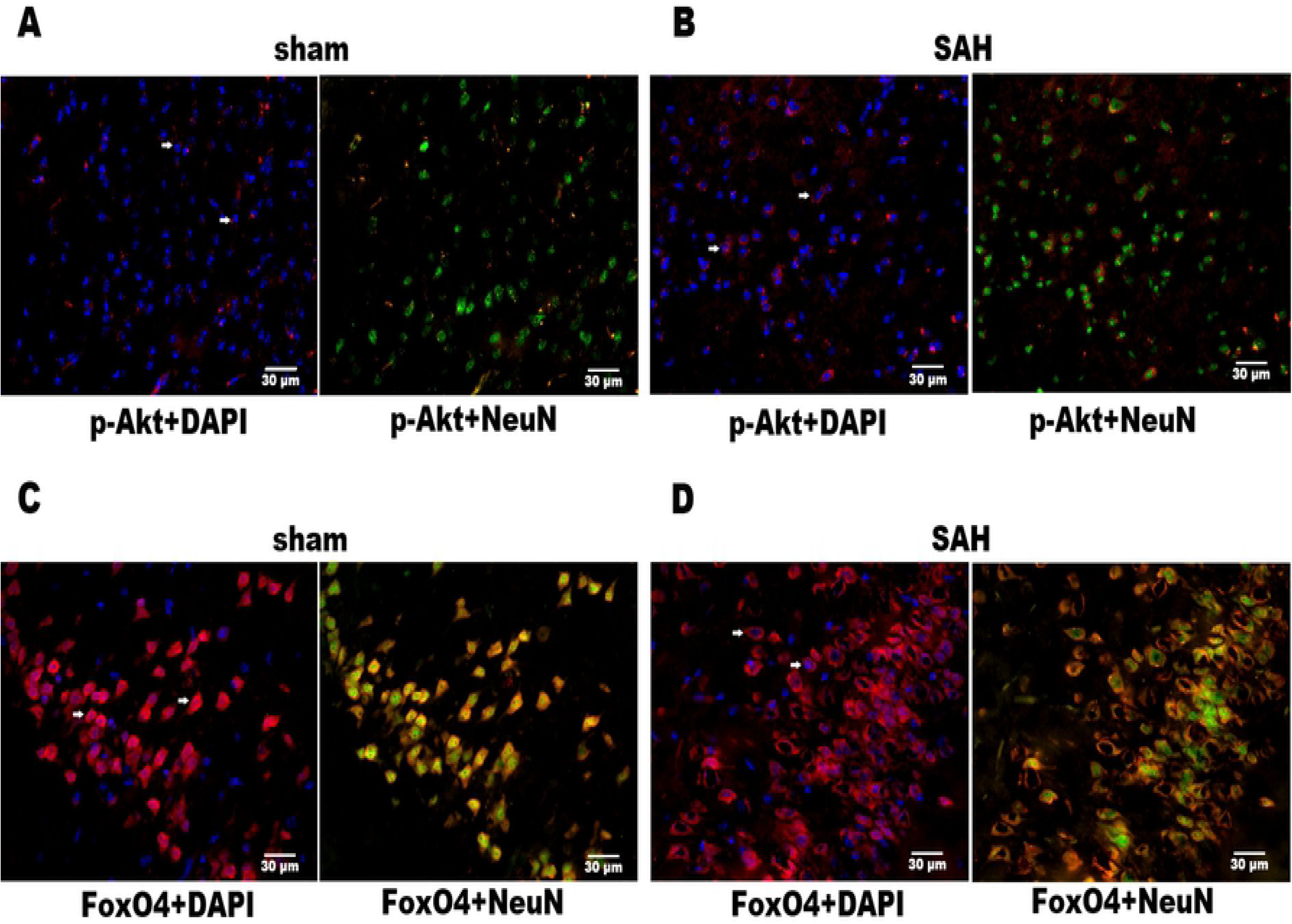
The changes of p-Akt and FoxO4 after SAH were detected qualitatively. P-Akt was detected in control group and SAH 1h group. The red fluorescence labeled p-Akt, green fluorescence labeled neurons, and DAPI labeled nucleus (A, B). FoxO4 was detected in control group and SAH group. FoxO4 was labeled by red fluorescence, neurons by green fluorescence, and nucleus by DAPI (C, D).

### 2.2. The change of the expression of FoxO4 after inhibition of Akt in vivo

After treatment with Akt inhibitor LY294002, supressed expression of p-Akt was confirmed(Fig.3A, F). We did not find the significant difference of FoxO4 in the total protein between the SAH group and LY294002 treatment SAH group(Fig. 3A, D). Nevertheless, the levels of FoxO4 protein in the nucleus were higher in the LY294002 treatment SAH group compared with the paired SAH group(Fig. 3B, E). This finding implies that inhibition of p-Akt could induce the nuclear translocation of FoxO4. The Cleaved-Caspase 3 was also detected in this study to indicate the apoptosis. We found Cleaved-Caspase 3 was also increased after LY294002 treatment.

**Figure 3.**
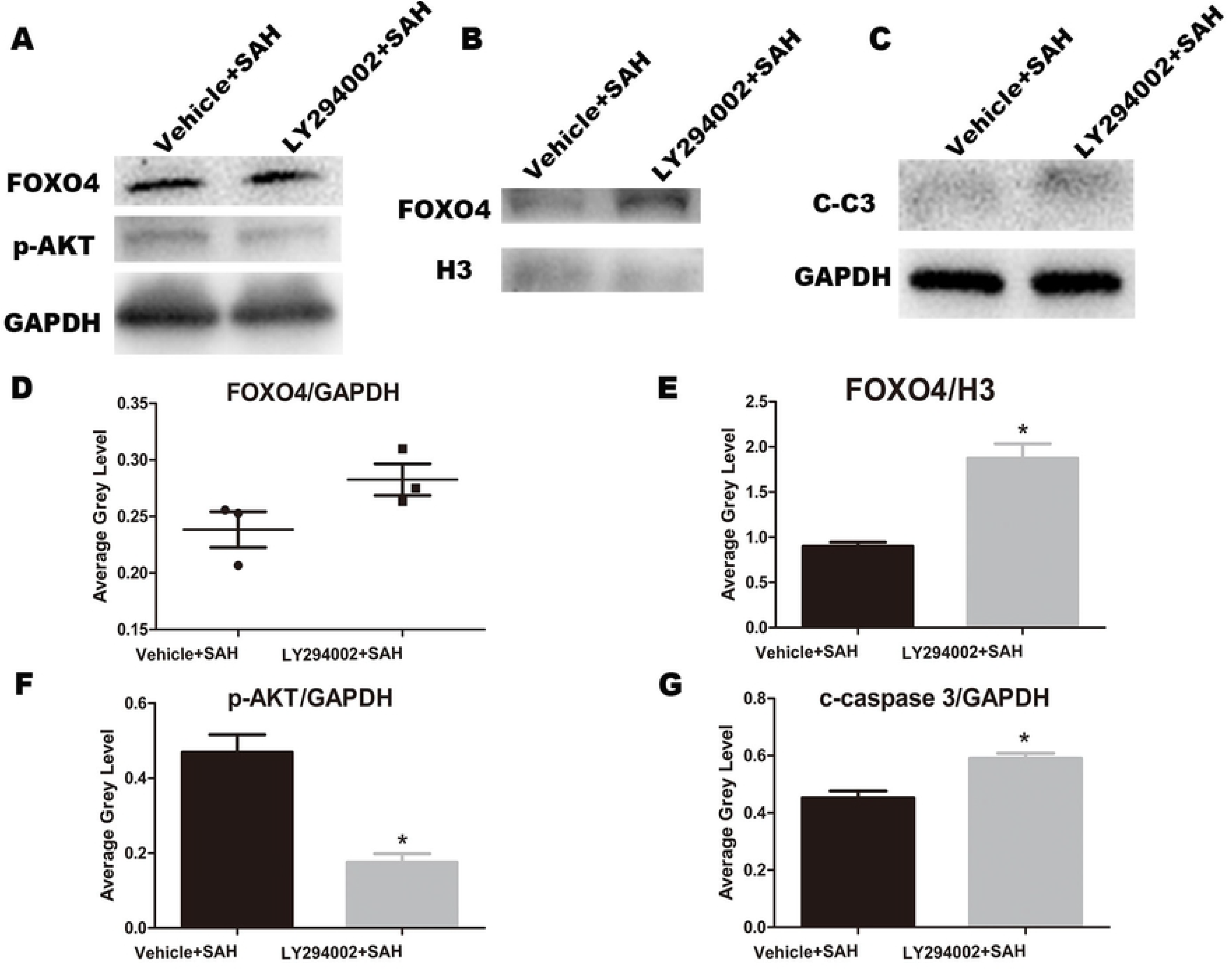
The changes after treatment with Akt inhibitor LY294002 *in vivo*. The total protein expressions of FoxO4 and p-Akt in group LY294002+SAH and group Vehicle+SAH were compared (A). The changes of FoxO4 nucleoprotein in group LY294002+SAH and group Vehicle+SAH were compared (B). We compared the changes of Cleaved-Caspase 3 between LY294002 + SAH group and group Vehicle+SAH group (C). Statistical analysis of FoxO4 protein compared between LY294002 + SAH group and group Vehicle+SAH group, n = 3, *P* > 0.05 (D). Statistical analysis of FoxO4 nuclear protein changes compared between LY294002 + SAH group and group Vehicle+SAH group, n = 3, \**P* < 0.05 (E). Statistical analysis of changes in p-Akt was compared between LY294002 + SAH group and group Vehicle+SAH group, n = 3, \**P* < 0.05 (F). We statistically analyzed the expression of Cleaved-Caspase 3 was compared between LY294002 + SAH group and group Vehicle+SAH group, n = 3, \**P* < 0.05 (G).

### 2.3. The effects of p-Akt inhibition on FoxO4 activation in neurons incubated with hemolyse in vitro

First, we detected the expression of p-Akt and FoxO4 in neurons incubated with hemolyse. The results showed that p-Akt was increased at 1h and 12h after the neurons were treated byh hemolyse(Fig. 4A, B). The expression of FoxO4 was increaed from 6h to 24h(Fig. 4A, C). After LY294002 treatment, p-Akt was suppressed sufficiently in this experimental system(Fig. 5A, E). the levels of FoxO4 in the total protein were not changed(Fig. 5A, D), but which in the nucleus were increased(Fig. 5B, F). This finding indicated the nuclear translocation of FoxO4. In our influence staining experiments, it was confirmed that FoxO4 was translocated to the nucleus after inhibition of p-Akt(Fig. 6A, B). The molecules related to apoptosis of neurons, Cleaved-Caspase 3, Bcl-2, and SOD2 were also investigated in the present studies. It was detected that Cleaved-Caspase 3 and SOD2 was increased significantly after treatment with LY294002. Bcl-2, an anti-apoptosis factor, was reduced after incubation of LY294002.

**Figure 4.**
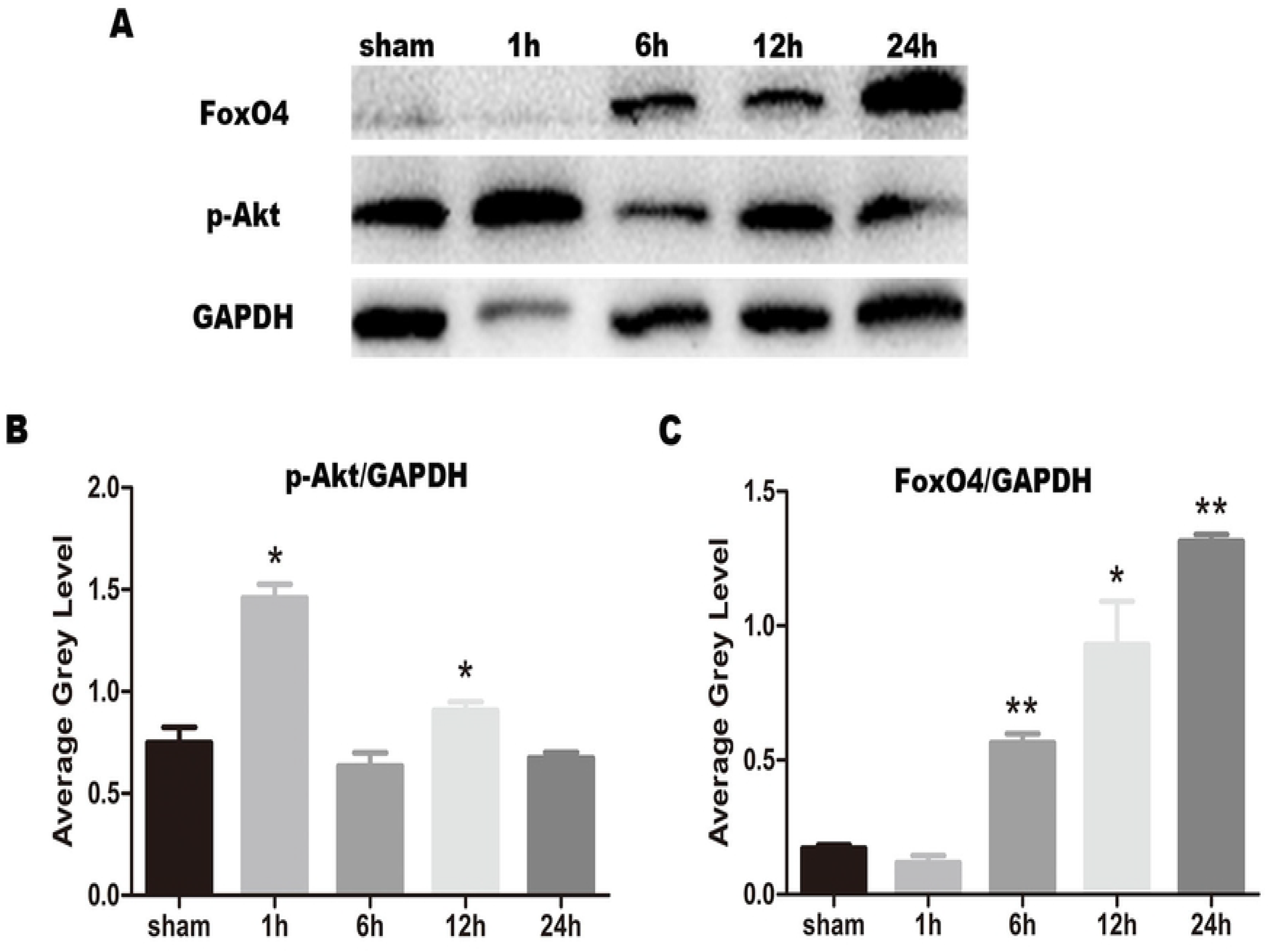
Expression of p-Akt and FoxO4 in neurons *in vitro*. Western blotting was used to detect the expression of FoxO4 and p-Akt *in vitro* (A). Statistical analysis of the expression of p-Akt in different time periods, n=3, \**P* < 0.05 vs. sham (B). The total protein expression of FoxO4 in different time periods was statistically analyzed, n=3, \**P* < 0.05 vs. sham, \*\**P* < 0.01 vs. sham(C).

**Figure 5.**
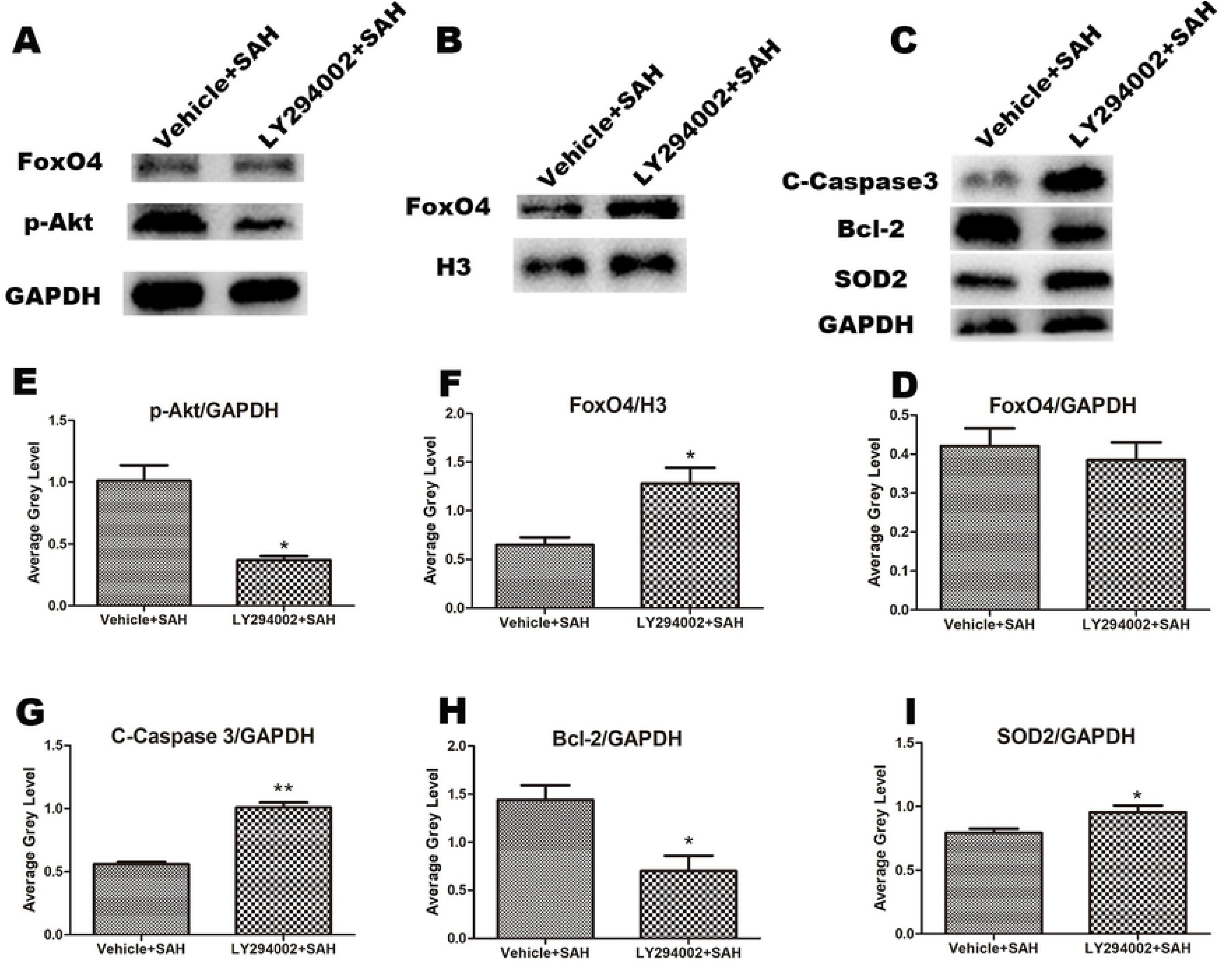
The changes after treatment with LY294002 were detected *in vitro*. Changes in FoxO4 and p-Akt in group LY294002+SAH and group Vehicle+SAH in total protein (A). Western blotting was used to detect the change of FoxO4 of nucleoprotein in group LY294002 + SAH and group vehicle + SAH (B). Cleaved-Caspase 3, Bcl-2 and SOD2 in group LY294002 + SAH and group vehicle + SAH were detected(C). The FoxO4 total protein in group LY294002+SAH and group Vehicle+SAH was analyzed statistically, n=3, *P* > 0.05 vs.vehicle + SAH (D). Statistical analysis of p-Akt in group LY294002 + SAH and group vehicle + SAH was detected, n = 3, \**P* < 0.05 vs. vehicle + SAH (E). The change of FoxO4 nucleoprotein in group LY294002 + SAH and group vehicle + SAH was statistically analyzed, n = 3, \**P* < 0.05 vs. vehicle + SAH(F). The changes of Cleaved-Caspase 3 in group LY294002 + SAH and group vehicle + SAH were statistically analyzed, n = 3, \*\**P* < 0.01 vs. vehicle + SAH (G). The changes of Bcl-2 in group LY294002 + SAH and group vehicle + SAH were statistically analyzed, n = 3,\**P* < 0.05 vs. vehicle + SAH (Figure 5H). The changes of SOD2 in group LY294002 + SAH and group vehicle + SAH were statistically analyzed, n = 3, \**P* < 0.05 vs. vehicle + SAH (I).

**Figure 6.**
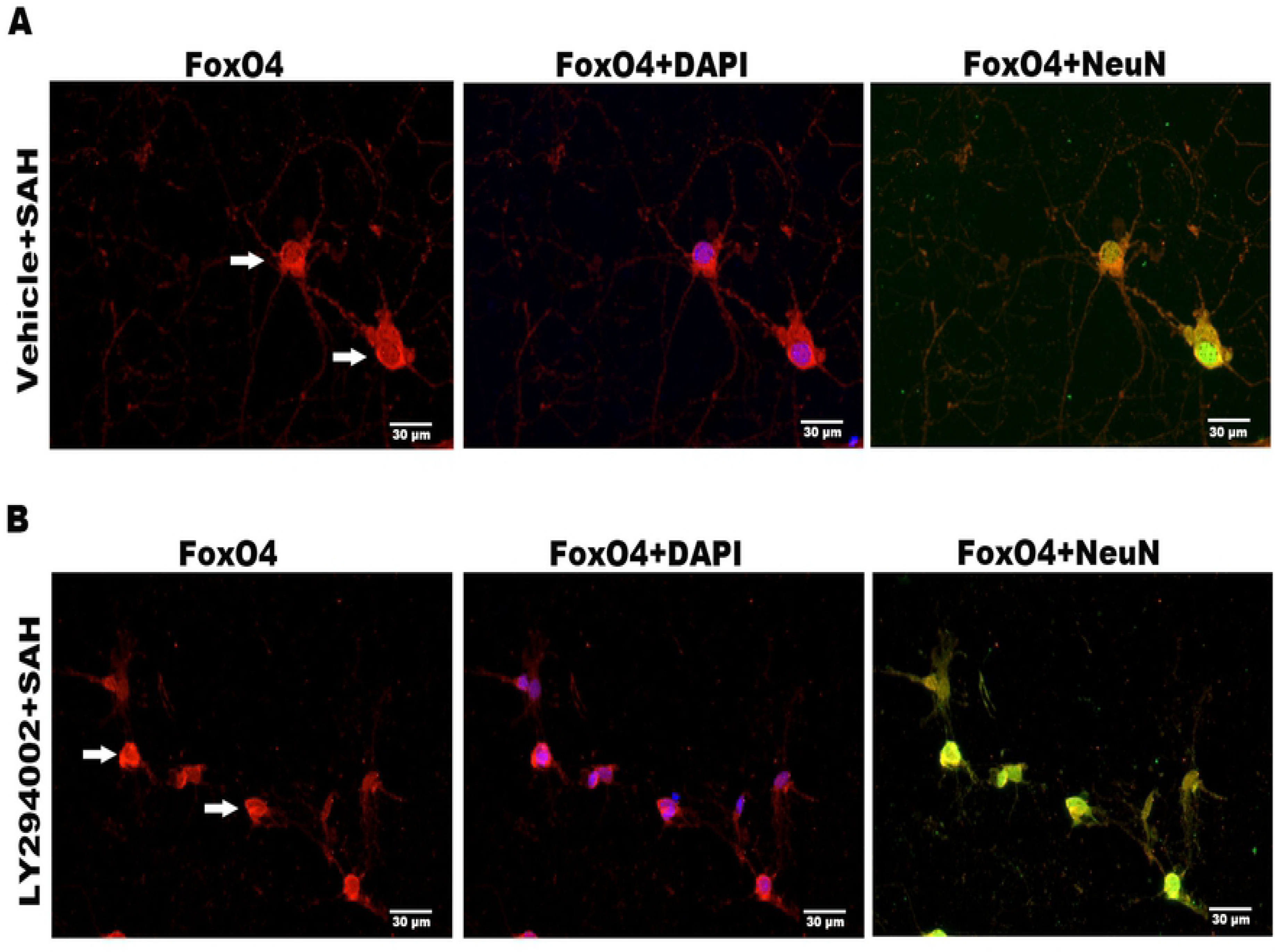
The distribution of FoxO4 in the cells was detected by immunofluorescence. FoxO4 in Vehicle+SAH group was located in cytoplasm, in which FoxO4 was labeled with red fluorescence, green fluorescence labeled neurons, and DAPI labeled nucleus (A). FoxO4 in the LY294002+SAH group was located in the nucleus, in which the red fluorescence labeled FoxO4, green fluorescence labeled neurons, and DAPI fluorescence labeled nucleus (B).

## 3. Discussion

The main results of this study showed that: 1) expression of FoxO4 in nucleus was increased from 6h to 24h after SAH after expreimental SAH model. This increase was confirmed in cultured neurons; 2) inhibiting Akt aggravated nuclear translocation of FoxO4 and may induce more apoptosis after SAH *in vivo* and *in vitro*.

FoxO4 is one member of FoxO family, which has been documented to be translocated into the nucleus in neurons after cerebral ischemia[11]. There are no studies focusing on the relationship between FoxO4 and SAH. Hence, as the first step, we detected the cytoplasmic and nuclear expressions of FoxO4 in the present study. We found nuclear FoxO4 was increased from 6h to 24h after SAH, indicating the activation of FoxO4 after SAH. As a surprising, both the cytoplasmic and nuclear expressions of FoxO4 were found to be increased at 12h and 24h. With regard to the levels of FoxO4 in total protein, no signficant changes were found either at 12h and 24h after SAH when compared to the sham group. We checked this result with additional animals, but the same results was got. The reason might be due to methods or the special characteristics of FoxO4. We are keeping trying to explore the reason in our laboratory.

FoxO4 and the other members in FoxO family are direct downstream targets of Akt. After dephosphorylation of Akt (inactive state), FoxOs translocates to nucleus where they bind DNA and regulate the genes related to metabolism, apoptosis, and reactive oxygen species (ROS) activation. We found the activation of both Akt (upregulation of p-Akt) and FoxO4 (translocation to necleus) after SAH. Endo er al have reported the increase of the p-Akt, which is consistent with our data. The changes of FoxO4 after SAH was investigated for the first time in the present study. Interestingly, the nuclear translocation of FoxO4 and the activation of Akt happened at the same time would be somewhat contradictory. We hypothesized that the increase in p-Akt is hemostatic responses to the activation of FoxO4 or some other downsteam targets. Hence, LY294002 was used in the present study to inhibit the increased p-Akt after SAH. After inhibitor of Akt, the FoxO4 increased even higher than the vehicle group both and *in vitro*. This finding indicated that the activation of Akt after SAH suppressed the activated FoxO4 to a higher level.

We also detected the Cleaved-Caspase 3 and Bcl-2 and SOD2 levels after LY294002 treatment *in vitro*. The Cleaved-Caspase 3, a key excutioner in apoptosis, was increased in LY294002 treatment group. On the other side, the Bcl-2, an antiapoptotic protein, was decreased after Akt inhibition. These finding implies that Akt/FoxO4 may induce apoptosis after SAH. SOD2 is the key enzymes that catalyse the detoxification of superoxide into oxygen and hydrogen peroxide, which is then converted to oxygen and water by catalase. In other words, the main biological function of SOD2 is to removes ROS in cells. The increase of SOD2 after Akt inhibition means the release of the function of FoxO4 to induce ROS production.

## 4. Materials and Methods

### 4.1. In vivo experiments

The Animal Care and Use Committee of Jinling Hospital approved the use of animals for this study, which was conducted in accordance with the Guide for the Care and Use of Labomiceory Animals published by the National Institutes of Health. Adult male C57/6JBL(30-35g) were purchased from the Animal Center of Jinling Hospital (Nanjing, China). The mice were housed in temperature-controlled and humidity-controlled animal quarters under a 12h light/12h dark cycle at 25°C, and provided with free access to food and water.

#### 4.1.1. *In vivo* subarachnoid hemorrhage(SAH) model

The mice were anesthetized with 10% chloral hydmicee (0.04mL/10g) and the SAH model was produced by stereotaxic insertion of a needle into the prechiasmatic cistern. Through a midline incision, the skin covering the anterior skull is opened. With a 0.9mm drill bit and a tail angle of 40°, a burr hole was drilled on the skull 4.5mm in front of the brainstem. The blood (50μL) was taken from C57/6JBL donors and passed through the burr hole at an angle of 40°with No.27 needle for 10 seconds until it reached the skull base. When the needle tip reached the base of the skull, it was retracted 0.5 mm. Then,Arterial blood was slowly injected into the prechiasmatic cistern over 20s under aseptic conditions. Stay in place for 5 min to avoid backflowand then after 30 min returned to their cages at room temperature.

#### 4.1.2. Experimental grouping and drug administmiceion

In the first experiment, 42 mice were assigned randomly to 7 groups: the control group (n = 6) and 6 SAH groups (1h, 6h, 12h, 24h, 3d and 5d, n = 6 for each group). The animals in the SAH 1h, 6h, 12h, 24h, 3d and 5d groups were subjected to experimental SAH and were killed at 1h, 6h, 12h, 24h, 3d and 5d after blood injection. The control mice were euthanized immediately after saline injection. Of the six samples in each group, three were used to extract total protein, and the other three were used to extract nuclear protein and cytoplasmic protein.

In the second experiment, 24 mice required. Akt inhibitor LY294002 was administered to investigate the significance of Akt in the nuclear translocation of FoxO4 after SAH. The mice were arranged in two groups, including SAH + vehicle (dimethyl sulfoxide [DMSO], Sigma, St. Louis, MO, USA) (25% DMSO in PBS; n = 8), SAH + LY294002 (50 mmol/L in 25% DMSO in PBS; n = 8). LY294002 (Sigma, St. Louis, MO, USA), and injected 10μL into the lateral ventricle (bregma: −0.5 mm, lateral: 1.0 mm, depth: 2.0 mm) at a mouse of 1μL/min as described in the previous studies[12]. SAH was performed 30 min after injection of medicine into lateral ventricle. In the experimental group (SAH + LY294002), 4 mice were euthanized at 1h to detect p-Akt, and 8 mice were euthanized at 24h (4 mice were tested for FoxO4 nuclear protein, and the other 4 mice were tested for FoxO4 whole protein). Similarly, in the control group (SAH + vehicle), we collected the whole protein of 1h and 24h and nucleoprotein of 24h, respectively, to detect the whole protein of p-Akt, FoxO4 and the nucleoprotein content of FoxO4.

#### 4.1.3. Western blotting

Each mouse was perfused with 150mL of ice-cold 0.9% saline through the left ventricle under deep anesthesia and brains were frozen until used. 1) **Total protein extraction:** Cortical tissues were lysed in RIPA buffer containing 1% PMSF and 1% phosphatase inhibitor, and protein concentration was measured by BCA method. 2) **Extraction of nucleoprotein and cytoplasmic protein:** Follow the instructions of nuclear protein and Cytoplasmic Protein Extraction Kit (Beyotime, Nantong, China). Under the condition of low osmotic pressure, the cells were fully expanded, then the cell membrane was destroyed, the plasma protein was released, and then the nucleus was precipitated by centrifugation. Finally, the nucleoprotein was extracted by high salt extraction reagent. Protein concentmiceions were measured using a detergent-compatible protein assay (Beyotime, Nantong, China).

Protein extracted by SDS-PAGE was electrotransferred to PVDF membrane. The membrane was sealed with 5% skim milk and incubated with primary antibodies against FoxO4(1:1000), p-Akt(1:1000), Cleaved-Caspase 3(1:1000) and SOD2(1:1000), with GAPDH (1:5000) and H3 (1:2000) as a loading control. Primary antibodies against FoxO4, H3 and GAPDH were purchased from Proteintech Group(Chicago, IL, USA); Primary antibodies against p-Akt, SOD2 and apoptosis-related proteins Cleaved-Caspase 3 were purchased from Cell Signaling Technology (Beverly, MA, USA). After the membranes were washed three times for 10 minutes each in PBST, they were incubated in the appropriate HRP-conjugated secondary antibody (1:400) for 2h. The blotted protein bands were visualized by enhanced chemiluminutesescence (ECL) Western blotting detection reagents (Amersham, Arlington Heights, IL, USA) and were exposed to X-ray film. Developed films were digitized using an Epson Perfection 2480 scanner (Seiko Corp, Nagano, Japan). Optical densities were obtained using Glyko Bandscan software (Glyko, Novato,CA, USA). All experiments were repeated at least three times.

#### 4.1.4. Immunofluorescence

The mice were perfused with 60mL of ice-cold 0.9% saline followed by 60mL of 4% formalin through the left ventricle under deep anesthesia. The brains were placed in 4% paraformaldehyde for 12 hours, after it was removed. Followed by 20%, 30% sucrose gradient dehydrated 1 day, until the brain completely sink to the bottom.Take out the brain tissue and let it dry on the surface, put it into the mold and bury it. After dewaxing, antigen repair, and cell lysis, the slides were incubated with anti-p-Akt (Cell Signaling Technology, Beverly, MA, USA; 1:1000) or anti-FoxO4 (Proteintech Group, Chicago, IL, USA; 1:2000) antibody overnight at 4°C. After PBS was removed, NeuN (EMD millipore, Billerica, MA; 1:200) was added 16-18h overnight. Then DAPI dye solution was added (Nanjing Kaiji Biological Co., Ltd.) and incubated at room temperature for 10 min. After washing with PBS, the slides were incubated with goat anti-rabbit IgG (diluted 1:500; Santa Cruz Biotechnology, Santa Cruz, CA, USA) and horseradish peroxidase (HRP) for 60 min at room temperature. The slices were washed three times with PBS (pH 7.4) for 5 min each. The slices were then slightly dried and sealed with anti-fluorescence quenching. The sections were observed under a fluorescence microscope and the images were collected.

### 4.2. In vitro experiments

#### 4.2.1. Culture and treatment of primary neurons

At 16-18 days of gestation, the mice were killed with neck broken and immersed in a culture dish containing 75% alcohol. After autoclave, the ophthalmic scissors opened the abdomen and took out the fetal mice. The head of fetal mice was severed and slightly washed in HBSS. We stripped the meningeal vessels and other tissues, removed the hippocampus and other brain parenchyma, and preserved the cortex. The cortex was cut up and a small amount of pancreatin was added, and then transferred into a 15mL centrifuge tube and bathed in 37°C water for 5 min. Added high glucose mediu (DMEM with 10% fetal bovine serum and 1% double antibody): pancreatin = 1.5:1, blowed gently and repeatedly. Centrifuged for 5 min at 1500 rpm, discarded the supernatant, and then suspended it again to obtain neurons with high purity, which could plant in the culture dish or 6-well plate coated with ploy-D. All the cells were replaced with the whole neuron culture medium. The proportion was neurobasal medium 100mL, B27 2mL, glumax 1mL, HEPES 750μL, Penicillin-Streptomycin (Liquid) 125μL (all from Gibco BRL, Grand Island, NY, USA). Two days later, the half fluid was changed (50%, all of which affected the growth of neurons). After 7 days, the cell protein or drug could be extracted for further treatment. In order to prevent cell pollution, all operations were completed under sterile conditions.

#### 4.2.2. *In vitro* subarachnoid hemorrhage(SAH) model

After 7 days of culturing primary neurons in whole neuron culture medium, the treatment can be started. Hemoglobin (sigma, St. Louis, Mo, USA) was diluted to 65mg/mL. hemoglobin was diluted with neurobasic medium, and then filtered with sterile filter. Discard the culture medium of neurons and replace it with half volume of hemoglobin (1mL for 6-well plate).The SAH model can be established *in vitro*.

#### 4.2.3. Experimental grouping and drug administmiceion

In the first experiment, The 30 well neurons cultured in the six well plate were randomly divided into 5 groups: the control group (n = 6) and four SAH groups (1h, 6 h, 12h, and 24h, n = 6 for each group). The SAH groups (1h, 6h, 12h, and 24h) were subjected to experimental SAH and the protein has been extracted at 1h, 6h, 12h and 24 h after Hemoglobin cultured. The neurons in the control group extracted protein directly after discarding the whole culture medium.

In the second experiment, the neurons were arranged in two groups, including SAH + vehicle (DMSO, Sigma, St. Louis, MO, USA) (1% DMSO in PBS; n = 18), SAH + LY294002 (20μmol/L in 1% DMSO in PBS; n = 18). LY294002 (Sigma, St. Louis, MO, USA). A 45 min pre-treatment with the treatment of SAH as described in the previous studies[13]. SAH induced after 45 min. When the establishment time of SAH *in vitro* reached one hour, the whole protein was extracted from the cells in six pores in both the experimental group(SAH + LY294002) and the control group(SAH + vehicle)(to detect p-Akt). In each group, six holes were used to extract total protein at the time of 24h after SAH (to detect FoxO4). The last six pores of each group were used to extract nucleoprotein in SAH 24 hours (to detect the expression of FoxO4 in nucleus).

#### 4.2.4. Western blotting

**1)Total protein extraction:** Total protein extraction: Primary neuron cell were lysed in RIPA buffer containing 1% PMSF and 1% phosphatase inhibitor, and protein concentration was measured by BCA method. **2)Extraction of nucleoprotein and cytoplasmic protein:** Follow the instructions of nuclear protein and Cytoplasmic Protein Extraction Kit (Beyotime, Nantong, China). Under the condition of low osmotic pressure, the cells were fully expanded, then the cell membrane was destroyed, the plasma protein was released, and then the nucleus was precipitated by centrifugation. Finally, the nucleoprotein was extracted by high salt extraction reagent. Protein concentmiceions were measured using a detergent-compatible protein assay (Beyotime, Nantong, China). The steps of Western blotting after protein extraction are consistent with the experiments *in vivo*.

#### 4.2.5. Immunofluorescence

After discarding the culture medium, the neurons were fixed with 4% paraformaldehyde. Antigen repair, cell membrane drilling and sealing(all from Beyotime, Nantong, China) were carried out in sequence at room temperature. The sections were then incubated with the diluted anti-p-Akt (Cell Signaling Technology, Beverly, MA, USA; 1:1000), anti-FoxO4 (Proteintech Group, Chicago, IL, USA; 1:2000) antibody overnight at 4°C. After PBS was removed, NeuN (EMD Millipore, Billerica, MA;1:200) was added and incubated at 4°C for 16-18h. After washing with PBS, DAPI dye solution was added (Nanjing Kaiji Biological Co., Ltd.; 1:50000) and incubated at room temperature for 10 min. After washing with PBS, the slides were incubated with goat anti-rabbit IgG (diluted 1:500; Santa Cruz Biotechnology, Santa Cruz, CA, USA) and horseradish peroxidase (HRP) for 60 min at room temperature. The slices were washed three times with PBS (pH 7.4) for 5 min each. The slices were then slightly dried and sealed with anti-fluorescence quenching. The sections were observed under a fluorescence microscope and the images were collected.

For immunofluorescence staining, positive cells were identified, and analyzed with the help of an investigator who was blind to the experimental treatments.

### 4.3. Statistical analysis

Comparisons between different groups were performed by analysis of variance (ANOVA) followed by Tukey’s multiple comparisons test if a significant difference had been determined by ANOVA. A probability value of *P* < 0.05 was considered statistically significant.

## 5. Conclusions

The activation of FoxO4 and increased phosphorylated Akt were detected after SAH. FoxO4 nuclear translocation was suppressed by Akt and which may related to neuronal apoptosis.

## Acknowledgments

Conceptualization, M.L.Z. and L.R.Z.; methodology, J.Q.L. and B.Y.; formal analysis, M.L.Z; investigation, X.L.L.; data curation, Y.L.H. and X.M.Z.; writing—original draft preparation, M.Q.; writing—review and editing, M.Q. and S.Q.G.; supervision, M.L.Z.; funding acquisition, M.L.Z.

